# Proximity RNA labeling by APEX-Seq Reveals the Organization of Translation Initiation Complexes and Repressive RNA Granules

**DOI:** 10.1101/454066

**Authors:** Alejandro Padròn, Shintaro Iwasaki, Nicholas T. Ingolia

## Abstract

Diverse ribonucleoprotein complexes control messenger RNA processing, translation, and decay. Transcripts in these complexes localize to specific regions of the cell and can condense into non-membrane-bound structures such as stress granules. It has proven challenging to map the RNA composition of these large and dynamic structures, however. We therefore developed an RNA proximity labeling technique, APEX-Seq, which uses the ascorbate peroxidase APEX2 to probe the spatial organization of the transcriptome. We show that APEX-Seq can resolve the localization of RNAs within the cell and determine their enrichment or depletion near key RNA-binding proteins. Matching the spatial transcriptome, as revealed by APEX-Seq, with the spatial proteome determined by APEX-mass spectrometry (APEX-MS) provides new insights into the organization of translation initiation complexes on active mRNAs, as well as exposing unanticipated complexity in stress granule composition, and provides a powerful and general approach to explore the spatial environment of macromolecules.

## Introduction

Proximity labeling has emerged as a valuable approach for understanding patterns of protein interaction and localization within cells. Proximity labeling techniques rely on enzyme-catalyzed *in vivo* reactions that mark targets near the labeling enzyme — which is typically fused to a query protein — and enable later, *ex vivo* analysis. The labeling reaction occurs within living cells and often acts over tens of nanometers. Proximity labeling is thus particularly well suited to capture transient, dynamic, and heterogeneous structures, complementing biochemical purifications that rely on direct and stable interaction.

The most dramatic advances in proximity labeling involve protein biotinylation, through the use of enzymes that produce diffusible reactive intermediates ‐‐ either short-lived radicals^1–3^, or longer lived adenylate esters^4–8^. These protein labeling tools can only indirectly address the organization of DNA in chromatin and RNA in various nuclear and cytosolic granules^9,10^. Currently, DNA can be labeled enzymatically through the action of an adenosine methyltransferase, in the DamID technique^11^. TRIBE is a similar approach for labeling RNAs with an adenosine deaminase^12^. While these base-modifying enzymes have proven valuable, for example in mapping binding sites of the heterochromatin protein CBX1 (HP1β)^13^, direct enzymatic modification operates only within a short, defined distance from the query protein and suffers from steric restrictions and other technical limitations.

Direct RNA proximity labeling promises new insights into the dynamic behavior of RNA. Translating RNAs move dynamically through the cytosol and often localize to specific regions of the cell^14–17^. Inactive RNAs can be sequestered into protein-RNA granules through a process of liquid-liquid phase separation (LLPS)^18–20^. Stress granule formation dynamically alter macromolecular localization, and mutations that increase stress granule formation or limit stress granule clearance are implicated in neurodegenerative diseases^21–23^. Recent work has argued that stress granules comprise two distinct components: a stable “core” surrounded by a concentration dependent “shell”^24^, which contain distinct macromolecules.

One powerful and distinctive approach for proximity labeling employs an engineered ascorbate peroxidase enzyme (APEX2) to convert a cell-permeable biotin-tyramide substrate into a highly reactive free radical that labels aromatic amino acids in proteins within ~25 nanometers^25,26^. APEX labeling has already provided insight into the protein composition of stress granules^27^. Motivated by the realization that nucleotides are also amenable to free radical-based chemistry^28^, we developed an RNA proximity labeling technique, using APEX2 (APEX-Seq) as a way to probe the spatial organization of the transcriptome. We show that APEX-Seq can resolve the localization of RNAs within the cell and determine their enrichment or depletion near key RNA-binding proteins. We then apply APEX-Seq to identify the transcripts that localize to stress granules and find that their composition varies depending on the stress applied. Our results reveal unanticipated complexity in stress granule contents, which likely extends to other RNA granules, and provides a powerful tool to explore this phenomenon.

## Results

### RNA biotinylation by the APEX proximity labeling enzyme

We reasoned that the radical mechanism underlying APEX proximity labeling of proteins^29^ would lead to similar, proximity-dependent biotinylation of RNA as well (Fig. 1a). Indeed, we found that purified recombinant APEX2 enzyme biotinylated RNA *in vitro*, in a reaction that depended on both biotin-tyramide and hydrogen peroxide, and similar to tyramide labeling of DNA by horseradish peroxidase (HRP)^28^ (Fig. 1b). Importantly, the biotinylation signal that we detected is RNase sensitive (Fig. 1c), and thus reflects labeling of the RNA in the reaction.

**Figure 1.**
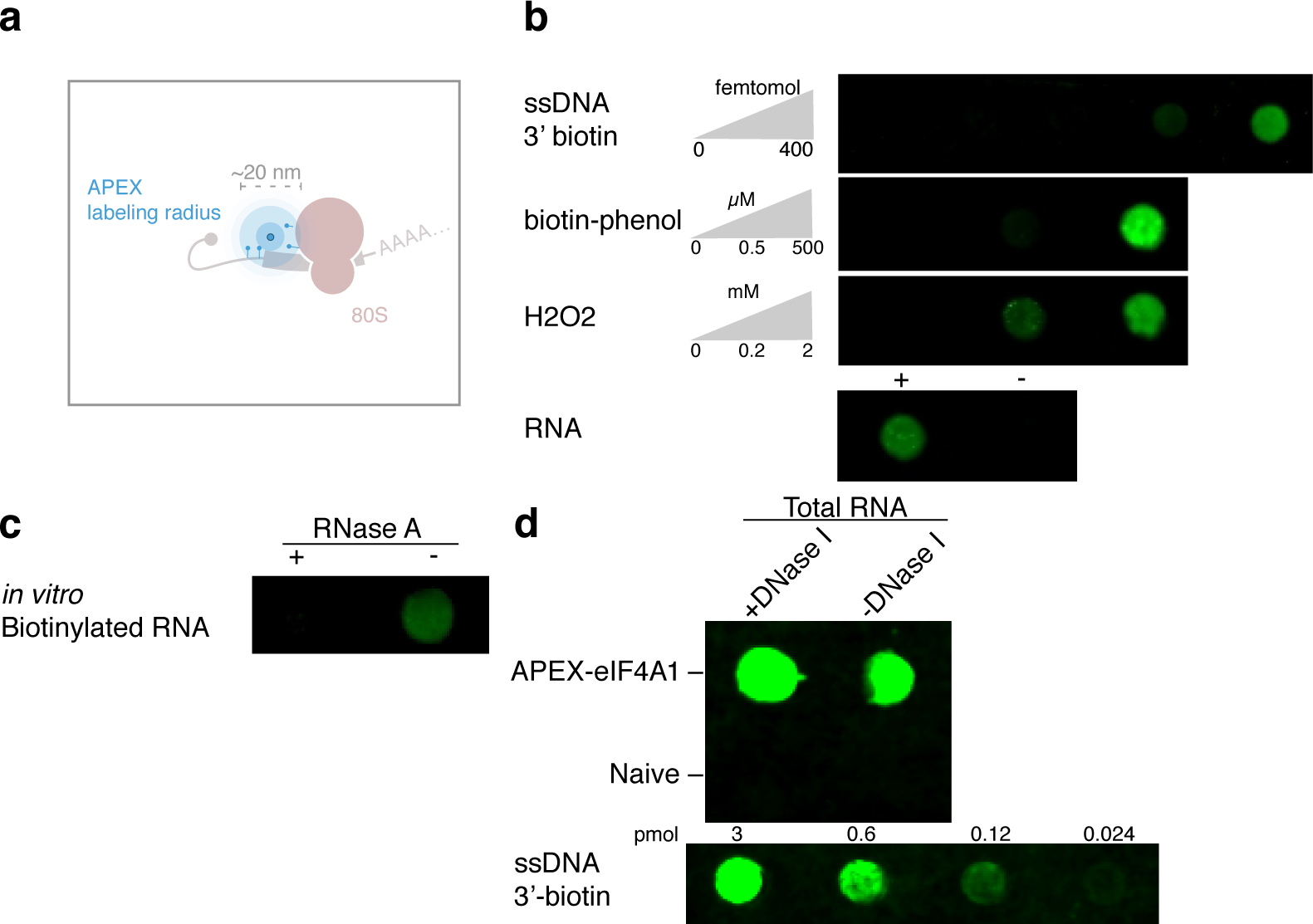
APEX proximity biotinylation of RNA. **a**, Diagram of APEX proximity biotinylation of both protein and RNA. **b**, *in vitro* labeling of RNA as a function of both biotin-phenol and hydrogen peroxide. **c**, *in vitro* biotinylated RNA subjected to RNase A treatment for 30 min at 37 °C. **d**, *in vivo* biotinylated RNA from an APEX2-eIF4A1 or Naive HEK293T cell line treated with DNase I for 30 min at 37 °C.

We next verified that the APEX labeling reaction biotinylates RNA *in vivo*. In order to test RNA biotinylation, we expressed APEX2 fused to the DEAD-box RNA helicase eIF4A1 in HEK293T cells (Supplementary Fig. 2a). This fusion retains its ability to bind RNA, including RocA-dependent stabilization on polypurine tracts^30^, which depend on the protein’s normal RNA-binding interface^30^ (Supplementary Fig. 1). The APEX2-eIF4A1 fusion also catalyzes protein biotinylation, leading to overall labeling roughly as strong as the labeling produced by APEX2-GFP, though with differences in the specific labeling pattern across the proteome (Supplementary Fig. 2b). Total RNA extracted from cells expressing this functional APEX2-eIF4A1 fusion is biotinylated after biotin-tyramide pre-incubation and peroxide treatment, in contrast to RNA from naïve cells treated in the same way, which shows no detectable biotin (Fig. 1d).

### APEX-Seq captures sub-cellular RNA localization patterns

Having shown that the APEX reaction biotinylates RNA as well as protein *in vivo,* we asked whether subcellular differences in RNA localization can be detected by purifying and sequencing this biotinylated RNA. We targeted APEX2 to three distinct locations within the cell: 1) the cytoplasm, through the use of APEX2-GFP; 2) the cytosolic face of the ER membrane, using a previously established C1(1-29)-APEX2 fusion^1^ containing the 29 N-terminal residues of cytochrome P450 2C1 (rabbit CYP2C1); and 3) the nucleus, using a CBX1-APEX2 fusion that links APEX2 to heterochromatin protein 1 beta (HP1β) (Fig. 2a). We carried out APEX labeling reactions in cells expressing each of these fusion proteins, purified the biotinylated RNA by streptavidin affinity, and analyzed both total and biotinylated RNA by deep sequencing. RNA-Seq read counts replicated extremely well in both total and streptavidin-purified samples (R^2^ ~ 0.99), confirming the reproducibility of our assay (Fig. 2b,c, Supplementary Fig. 3). Moreover, we saw distinct patterns of enrichment and depletion after purifying biotinylated RNA by streptavidin affinity, suggesting that different APEX fusion proteins were labeling different transcripts (Supplementary Fig. 4a).

**Figure 2.**
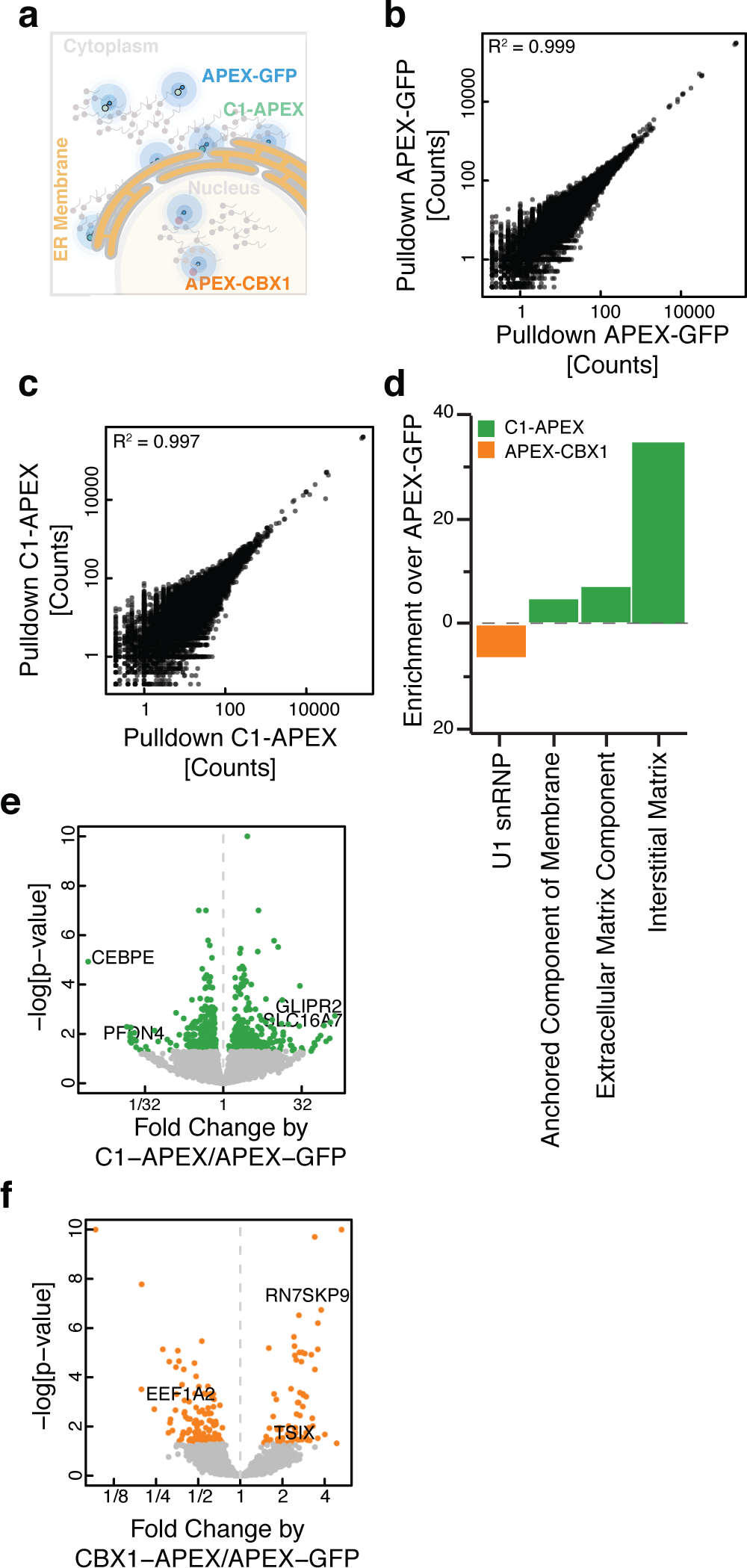
APEX proximity biotinylation reveals RNA sub-cellular localization. **a**, Diagram of localization using APEX-Seq. **b**, **c**, Correlation plot showing agreement between replicate samples for streptavidin affinity purification of biotinylated RNA from (**b**) an APEX2-GFP cell line and (**c**) streptavidin-pulldown C1-APEX2 cell line. **d**, GO term enrichment for transcripts preferentially labeled in C1-APEX2 and CBX1-APEX2 cell lines. **e**, **f**, volcano plot for (**e**) C1-APEX2 and (**f**) CBX1-APEX2 compared to APEX2-GFP showing enrichment and depletion for specific RNAs.

We next wanted to test whether these patterns of biotinylation reflected the sub-cellular localization of RNAs relative to the APEX fusion protein. We compared biotinylated RNA from C1-APEX2 and CBX1-APEX2 fusions against the RNA labeled by our diffuse APEX-GFP control. We found that C1-APEX2 labeling enriched strongly for mRNAs encoding membrane-associated proteins, reflecting their localization to the surface of the ER during co-translational secretion (Fig. 2d, Supplementary Fig. 3,4b). The highest enrichment scores we saw included 139x enrichment of Golgi-Associated Plant Pathogenesis-Related Protein 1 (GLIPR2) mRNA and 86x enrichment of the SLC16A7 transcript, which encodes a bidirectional transport of short-chain monocarboxylates^31^. In contrast, transcripts encoding soluble cytosolic proteins such as PFDN4, a subunit of the heterohexameric chaperone prefoldin, and nuclear proteins such as the bZIP transcription factor CEBPE, were depleted in C1-APEX2 relative to APEX2-GFP ^31,32^ (Fig. 2e, Supplementary Fig. 4d).

Likewise, we found that the nuclear CBX1-APEX2 fusion preferentially labeled non-coding nuclear RNAs (Fig. 2d). We saw a strong enrichment of the 7SK small nuclear pseudogene (RN7SKP9), as well as an enrichment of TSIX, the antisense transcript derived from the XIST locus. By contrast, we saw depletion of the mRNA encoding for Elongation Factor 1 Alpha 2, a highly abundant coding transcript (Fig. 2f). Taken together, the data from our C1-APEX2 and CBX1-APEX2 labeling show that APEX-Seq captures patterns of RNA localization across cellular compartments even when these are not separated by membranes.

### APEX-Seq captures protein-RNA interaction patterns

We next asked whether patterns of protein-RNA interaction, occurring on an even finer length scale, could be detected by APEX-Seq enrichment. In addition to the APEX2-eIF4A1 fusion described above, we fused APEX2 to the 7-methylguanosine (7mG)-cap binding protein eIF4E1 (Fig. 3a). We compared biotin-labeled RNAs from cells expressing APEX2-eIF4E1 against total RNA and observed the depletion of non-coding RNAs with 2,2,7-trimethylguanosine caps that bind eIF4E1 with far lower affinity^33^ (Supplementary Fig. 4c). Intriguingly, we observed the strongest enrichment for immediate early genes FOSB and EGR3 (Fig. 3b). Immediate-early genes are preferentially translated when the eIF4E1-interacting scaffold eIF4G is limiting^34^, and enrichment of these transcripts may reflect their preferential association with eIF4E1. Several proto-oncogenes, including c-Myc, are also preferentially translated in conditions of limiting eIF4G^34^. Indeed, eIF4E1 is an oncogene whose overexpression promotes cellular transformation^35,36^. In line with these observations, we find that APEX2-eIF4E1 preferentially labels the oncogenic C-MYC, VEGF-A, and FGF13 mRNAs (Fig. 3c). More broadly, protein synthesis of long, structured 5’ UTR-containing mRNAs appear to be sensitive to eIF4E1 levels^35,37,38^, and we find greater enrichment of long 5’-UTR containing mRNAs upon labeling by APEX2-eIF4E1 (Fig. 3d). The eIF4E1 transcript was also enriched, perhaps reflecting nascent APEX2-eIF4E1 biotinylating its own mRNA (Fig. 3b).

**Figure 3.**
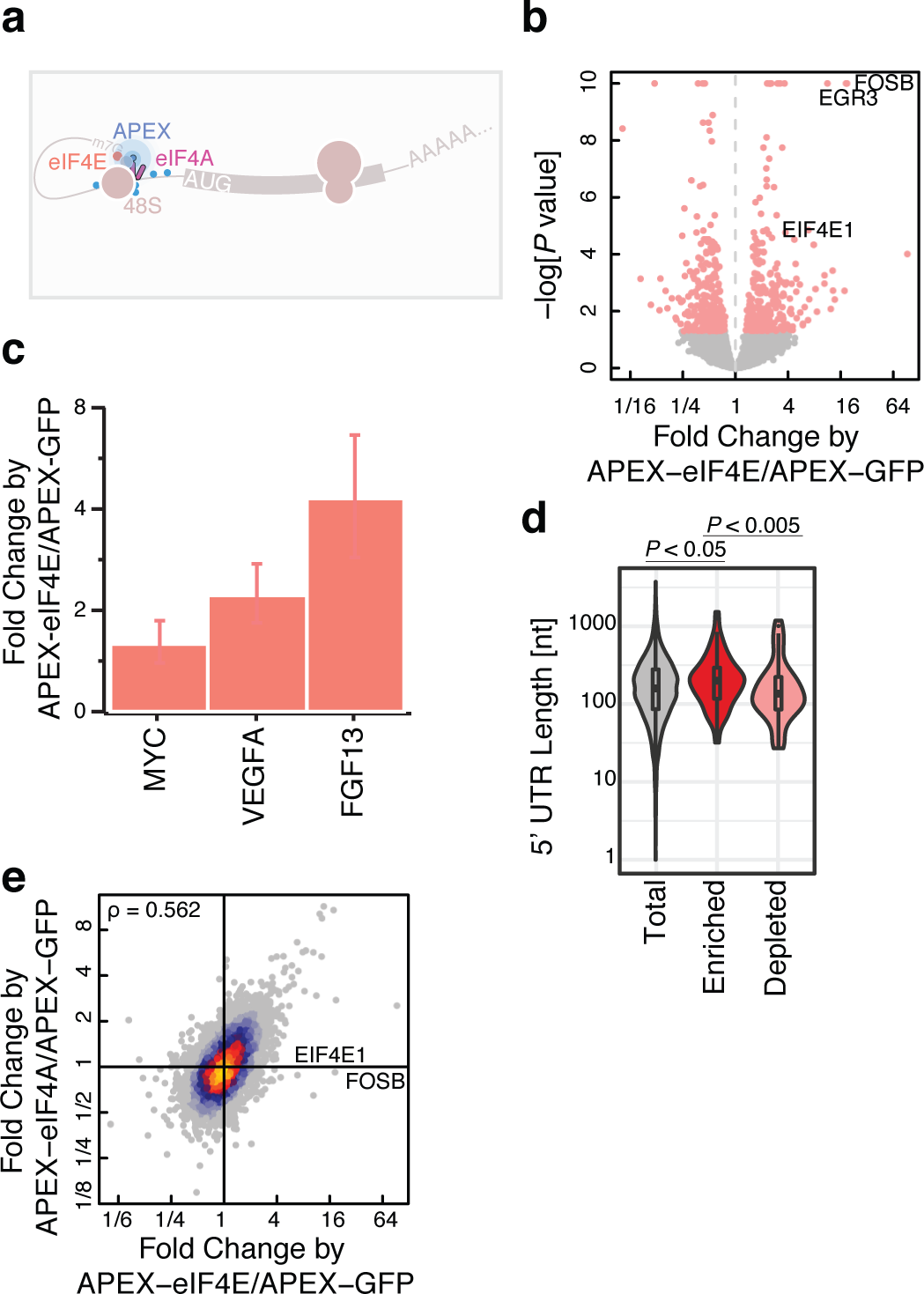
APEX proximity biotinylation reveals ribonucleoprotein complex composition. **a**, Diagram of the 43S preinitiation complex. Blue sticks signify biotin adducts formed from the proximity labeling reaction catalyzed by APEX2. **b**, Volcano plot for APEX2-eIF4E1 showing enrichment and depletion of specific RNAs compared to APEX2-GFP. **c**, Barplot showing enrichment for mRNAs encoding cell transforming proteins in APEX2-eIF4E1. **d**, 5’ UTR length distribution for APEX2-eIF4E1 enriched or depleted RNAs. P-values calculated using the Mann-Whitney test. Enrichment and depletion is calculated as statistically significant (FDR <0.05) and either top or bottom 2.5% of the fold change. **e**, Correlation plot between APEX2-eIF4E1 and APEX2-eIF4A1 datasets in comparison to APEX2-GFP, highlighting uniquely enriched mRNAs with respect to APEX2-eIF4E1.

Next, we asked whether APEX2-eIF4E1 and APEX2-eIF4A1 showed similar labeling patterns, since both bind to eIF4G as components of the eIF4F translation initiation complex. Indeed, we found good correspondence between their respective labeling patterns (Fig 3e). Both proteins enrich for mRNAs encoding translation elongation factors, which are known to show a particular sensitivity to inhibition of eIF4E1^39,40^ and eIF4A1^30^. While eIF4E1 and eIF4A1 labeling patterns overlapped substantially, they were not identical. For example, the mRNA encoding eIF4E1 is enriched only in APEX2-eIF4E1 labeling and not in APEX2-eIF4A1 labeling, consistent with the model that this enrichment reflects biotinylation by nascent protein. The similarity between eIF4E1 and eIF4A1 labeling patterns reflects the co-localization of these proteins, as we find a substantially weaker correlation in RNA labeling between CBX1-APEX2 and APEX2-eIF4A1 fusions (Supplementary Fig. S4e).

### Proximity proteomics places eIF4A1 on the 3’ side of the 48S preinitiation complex

Our observation that APEX can label RNA as well as protein now offers the ability to match spatially resolved transcriptomic and proteomic data. We therefore performed quantitative tandem mass tag (TMT) mass spectrometry^41^ in cells expressing APEX2-eIF4A1 (Supplementary Fig. 2). Ratiometric analysis using TMT labeling showed reproducible quantitation between biological replicates (Spearman’s ρ ~ 0.98; Supplementary Fig. 5a,b). We find that eIF4A1 labeling enriches other components of the eIF4F complex — the mRNA cap-binding protein eIF4E1 and the scaffolding protein eIF4G1 — as well as the poly(A) binding protein, PABPC1, which binds eIF4G1, and several small subunit ribosomal proteins (Fig. 4a,b). More broadly, APEX2-eIF4A1 spatial proteomics shows enrichment for translation initiation, RNP organization, and post transcriptional regulators in gene ontology enrichment analysis (Fig. 4c). This pattern of enrichment is similar to BioID analysis of eIF4A1, which relied on long duration (24 hour) labeling by an eIF4A1-birA* fusion protein. We saw a substantial (~25%) overlap between the proteins enriched in our rapid (<1 minute) APEX labeling experiment and those seen in long-term BioID labeling (Supplementary Fig. 5d; hypergeometric p < 0.01). Furthermore, we saw a substantial correlation in the quantitative pattern of labeling (Supplementary Fig. 5e; ρ = 0.241) despite the different chemical bases for APEX and BioID biotinylation.

**Figure 4.**
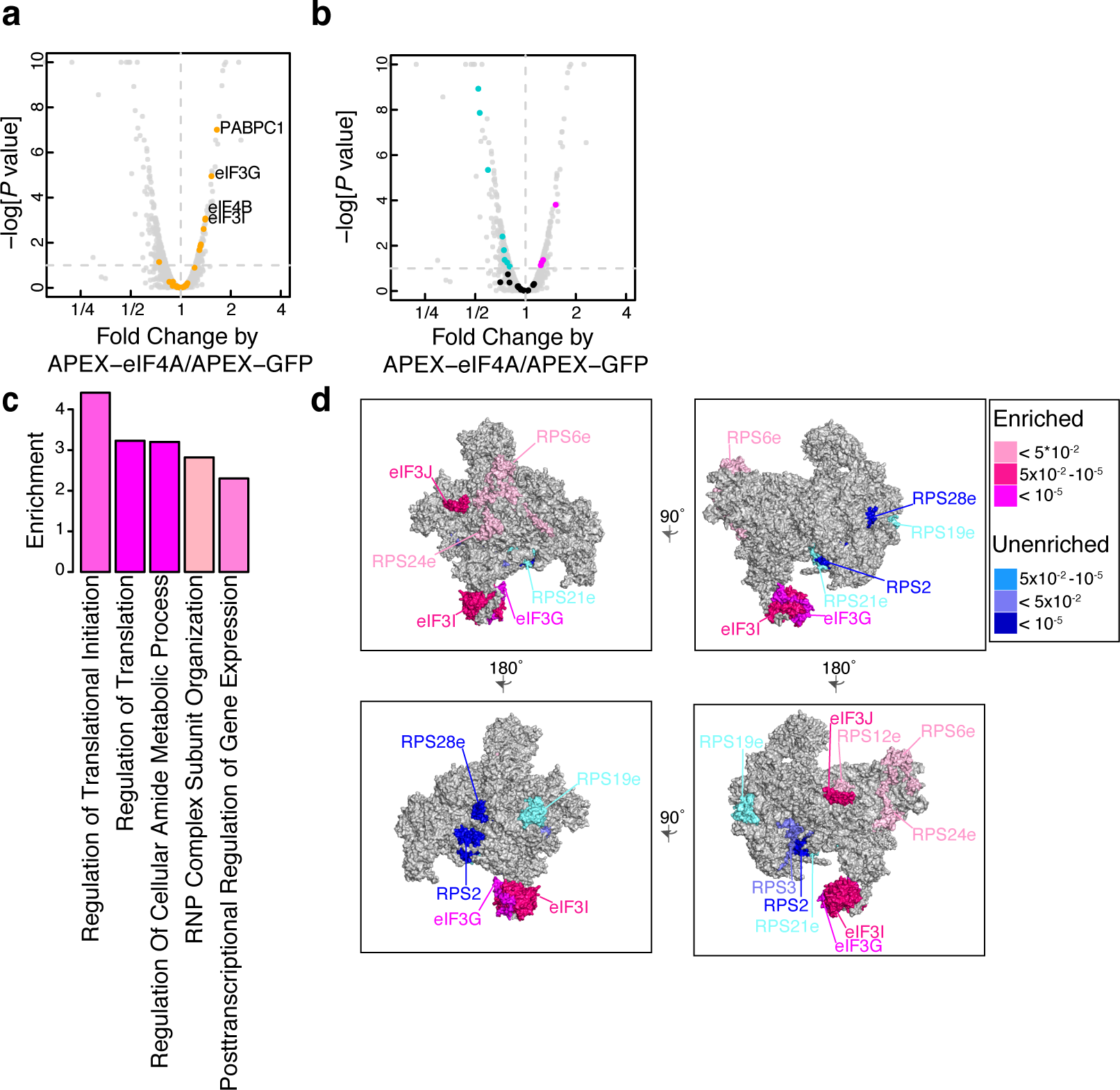
Proximity proteomics places eIF4A1 on the 3’ side of the 43S preinitiation complex. **a**, Enrichment of translation initiation factors (orange) in APEX2-eIF4A1 proximity labeling relative to APEX2-GFP. **b**, Enrichment and depletion of small subunit ribosomal proteins in APEX2-eIF4A1 proximity labeling. Significantly enriched proteins in pink, depleted in blue, and others in black. **c**, GO term enrichment for APEX2-eIF4A1 as compared to APEX2-GFP. **d**, Mapping enriched (pink) or depleted (blue) eIF3 and RPS subunits onto the eukaryotic pre-initiation complex. Different perspectives of the 43S complex are shown. Differently colored proteins signify FDR corrected p-values.

We noticed a striking pattern of enrichment for some small subunit ribosomal proteins and depletion of others, despite their uniform presence in 43S pre-initiation complexes (Fig. 4b, Supplementary Fig. 5c). To better understand why different 40S proteins show enrichment or depletion in APEX2-eIF4A1 proteomics, we mapped these proteins onto the structure of the eukaryotic 43S preinitiation complex^42^. We find that the proteins enriched by APEX2-eIF4A1 are situated near the mRNA entry site of the 43S preinitiation complex, on the side towards the 3’ end of the mRNA, while the depleted proteins lie closer to the mRNA exit site (Fig. 4d). The 43S preinitiation complex also binds to eIF3, a large, multi-protein complex that plays diverse roles in translation initiation. Enriched eIF3 subunits were also localized toward the mRNA entry site^43^. These data clearly place the RNA helicase eIF4A on the leading edge of the 43S complex, addressing an open question about the organization of this complex^44^.

### Proximity labeling captures the dynamic proteomic landscape during stress granule assembly

In addition to translation initiation factors, we noticed that APEX2-eIF4A1 labeling enriched a number of stress granule proteins (Supplementary Fig. 5f), although we saw no evidence of stress granule formation in these unstressed cells (Supplementary Fig. 6a). Our observation was consistent with previous studies reporting the presence of eIF4A1 in stress granule cores^24^, which are pre-existing structures present prior to SG formation^45^, as well as the BioID proximity labeling of stress granule components after long term (24 hour) expression of eIF4A1-birA* in unstressed cells^46^. APEX2-eIF4A1 labeling after stress resulted in an ~100x enrichment of biotin signal inside granules relative to the overall cytoplasm (Fig. 5a, Supplementary Fig. 6b), demonstrating a strong and specific enrichment of APEX2-eIF4A1 within stress granules after heat shock. These results suggested that we could use our APEX2-eIF4A1 fusion to study the stress granule transcriptome and proteome with short (<1 minute) proximity labeling, allowing us to track the dynamics of SG assembly.

**Figure 5.**
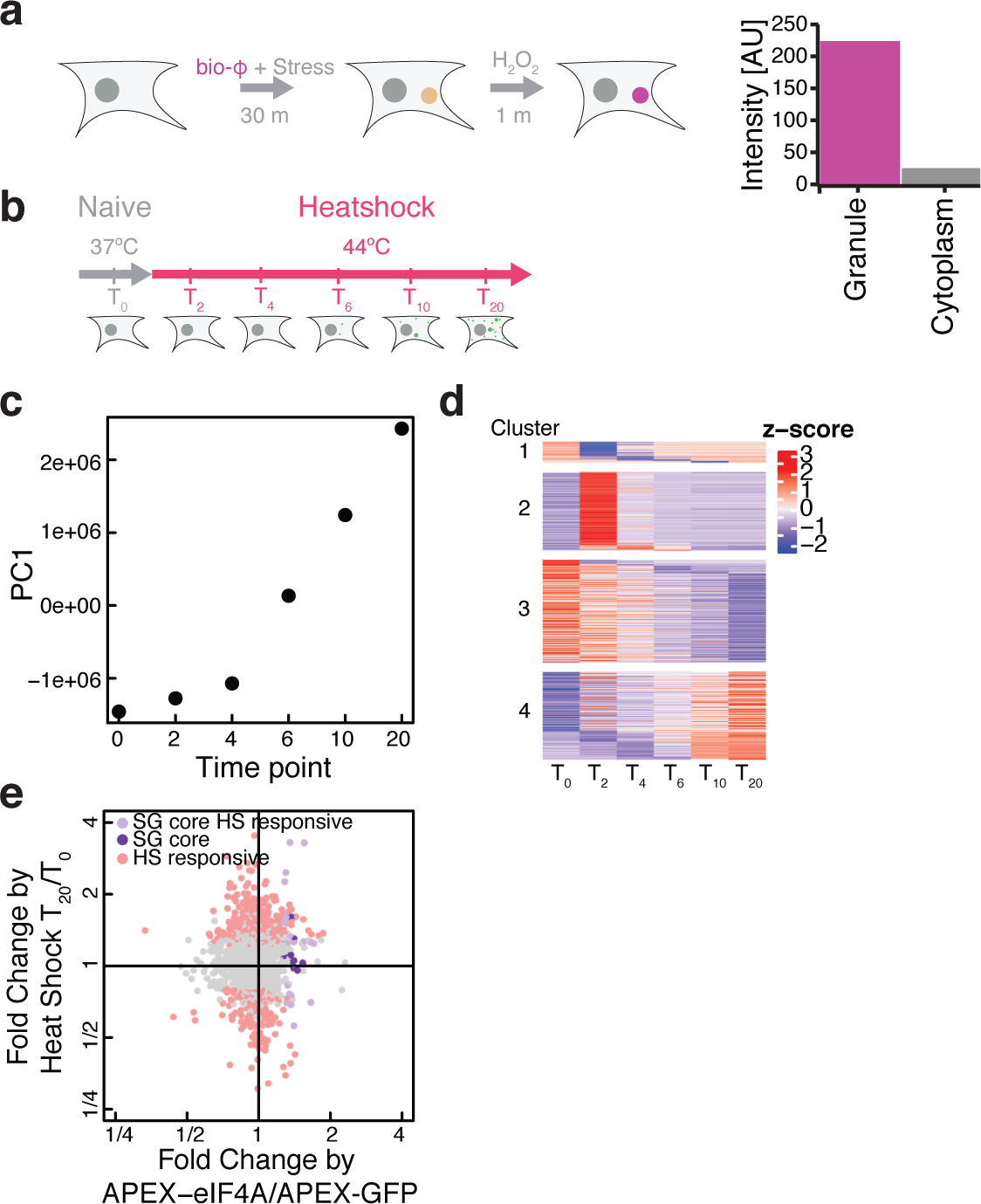
Timecourse of stress granule assembly following heat shock.

**a**, Diagram of APEX2-eIF4A1 immunofluorescence assay with thapsigargin treatment showing the biotinylation intensity of APEX2-eIF4A1 inside and outside stress granules. **b**, Diagram of APEX2-eIF4A1 heat shock time course. **c**, First principle component as a function of heat shock stress time points. **d**, Heatmap result from ImpulseDE2 highlighting significantly changing (FDR corrected p-value <0.01) proteins that: decrease transiently with respect to heat shock (Cluster 1), increase transiently over time (Cluster 2), decrease gradually over time (Cluster 3), or increase gradually over time (Cluster 4) with respect to heat shock. **e**, Comparison between T_20_ post heat shock with respect to pre heat shock (T_0_) vs APEX2-eIF4A1 with respect to APEX2-GFP. Heat shock responsive proteins (light pink, FDR corrected p-value < 0.05), stress granule core proteins that are heat shock insensitive (purple, FDR corrected p-value < 0.05), stress granule core proteins that are heat shock sensitive (lavender, FDR corrected p-value < 0.05).

We thus used APEX-Seq along with APEX-MS to measure the changing enrichment patterns of eIF4A1 labeling during the rapid and synchronous formation of stress granules following heat shock (Supplementary Fig. 7). We monitored the dynamics of stress granule assembly across a 20-minute time course following 44 °C heat shock (Fig. 5b). Replicates from our quantitative TMT-labeled proteomics correlated well (Supplementary Fig. 8a,b, Spearman’s ρ ~ 0.98). We observed substantial changes across the time course, and principal component analysis of these changes revealed that the first principal component captured ~50% of variation and described an increasing trend across the time course of SG assembly (Fig. 5c). The inflection point in this increase occurred after 5 minutes, which corresponds to the time when microscopically observable stress granules appear (Supplementary Fig. 7b). The component loadings from PC1 were dominated by increases in SG-related proteins like G3BP1 along with decreases in translation initiation and elongation factors (Supplementary Fig. 8c).

Hierarchical clustering segregated proteomics data from unstressed controls and early (2 - 4 minute) timepoints away from later (6 - 20 minute) samples (Supplementary Fig. 8d). This distinction between early and late timepoints also agrees with the timecourse of microscopically observable SG assembly (Supplementary Fig. 7b). We therefore partitioned our data into overlapping “early” (0 - 4 min), “middle” (2 - 10 min), and “late” (6 - 20 min) categories and identified proteins enriched in each of these groups. This analysis highlighted a number of important SG forming proteins in the “middle” timepoints, such as CAPRIN1^47,48^ (Supplementary Fig. 9). The stress granule forming protein, G3BP1, which interacts with CAPRIN1^48^, appeared in the “late” timepoints, suggesting an order of events in interactions with respect to eIF4A1. More broadly, GO terms for RNA localization are enriched during the late time point (Supplementary Fig. 8e).

We took advantage of our high time resolution and further refined this analysis by fitting the dynamics of eIF4A1 proximity labeling with impulse models using ImpulseDE2^49^ (Fig. 5d). This approach can identify proteins showing unidirectional trajectories of increased or decreased labeling, in addition to finding those showing transient changes. This analysis agrees with our hierarchical clustering results, as we find that “middle” timepoint proteins such as CAPRIN are induced early in the impulse model (cluster 2), whereas “late” proteins like G3BP1 more gradually accumulate over time (cluster 4). Overall, these data highlight the dynamic nature of the stress granule proteomic landscape as it is assembled upon heat shock.

The granule proteome at our final timepoint appeared broadly consistent with the proteins identified in previous proximity labeling studies, carried out after long stress induction regimes when granules were fully assembled. The quantitative changes in APEX2-eIF4A1 labeling that we saw after 20 minutes of heat shock correlated with the changes in BioID labeling induced by 3 hours of arsenite treatment (ρ = 0.24). This modest correlation was enhanced by restricting our analysis to proteins with an established eIF4A1 interaction or stress granule localization (ρ = 0.54), suggesting that these two very different labeling strategies captured the same underlying stress granule proteome. While BioID data suggest that stress granule formation is driven by pre-existing interactions that remain largely unchanged after stress^46^, we find a substantial and coherent change in the pattern of eIF4A1 proximity labeling across stress granule assembly. The rapid (< 1 minute) labeling enabled by APEX, in conjunction with our high-resolution timecourse, allowed us to resolve these dynamics more clearly.

We also saw a strong correlation between proteins showing stress-induced APEX2-eIF4A1 labeling in our study and previous proximity labeling studies using the granule marker G3BP1. This enrichment was strongest for our “middle” class of proteins (p < 1e-10, hypergeometric test), and was also significant for our “late” class (p < 0.02). These G3BP1-proximal proteins overall showed significantly higher labeling in our analysis of “middle” timepoints, relative to other proteins, as well (p < 1e-15, Wilcoxon test). Notably, we find G3BP1 in our “late”-enriched group and it shows heat shock responsive labeling by APEX2-eIF4A1; conversely, previous work found stress-dependent labeling of eIF4A1 by APEX-G3BP1. These results are consistent with G3BP1 and eIF4A1 co-localizing to stress granules and thus showing stronger mutual labeling and more similar overall labeling patterns after stress.

While stress granules are dynamic structures *in vivo*, previous work has identified a stable “core” that can be purified biochemically^24,50^. In order to better understand how proximity labeling patterns mapped onto stress granule assembly, we investigated these core stress granule proteins in our data. We defined a set of “stress responsive” proteins that were statistically enriched in APEX2-eIF4A1 labeling, in comparison with APEX2-GFP, and were further enriched in a comparison between APEX2-eIF4A1 labeling of the post-shock (T_20_) time point with respect to the pre-shock (T_0_) time point. In contrast, proteins labeled “stress insensitive” are those that were not significantly enriched or depleted in a comparison between APEX2-eI4FA1 labeling of the pre-shock (T_0_) and latest post-shock (T_20_) timepoints, but were enriched in a comparison between APEX2-eIF4A1 as compared to APEX2-GFP. Interestingly, we found a subset of the stress granule core components to show stress-insensitive eIF4A1 enrichment (Fig. 5e). Among these are ATXN2, a protein implicated in amyotrophic lateral sclerosis (ALS)^51^.

Previous work has implicated low complexity protein domains in stress granule formation^19^. We asked if there were other protein features predictive of stress granule residency. To do this, we compared our final post-shock time point (T_20_) with respect to our pre-shock time point (T_0_). We find the presence of RNA recognition motifs (RRMs) to be most predictive of RNA granule residency, along with an array of protein-protein interaction domains, whereas DEAD-box RNA helicases are depleted (Supplementary Fig. 10). Prion propensity^52^ does not predict SG residency in our analysis. These data suggest a central role for RNA interactions in recruiting proteins to stress granules and a role for some RNA helicases in opposing granule assembly. Perhaps translating ribosomes increase the likelihood that RNA remodeling enzymes associate and prevent interactions between the translating mRNAs and key RNA binding proteins.

### APEX-Seq reveals that different stresses yield distinct granule RNAs

We analyzed the SG transcriptome by performing APEX-Seq on cells subjected to heat shock as well as cells treated with hippuristanol, a drug that directly targets eIF4A1 and induces RNA granule formation through a phospho-eIF2α-independent mechanism^53^. While hippuristanol itself disrupts the eIF4A / RNA interaction, eIF4A1 is readily recruited into hippuristanol-induced stress granules^54^. Both heat shock and hippuristanol cause substantial changes in the pattern of APEX2-eIF4A1 labeling (Supplementary Fig. 11a,b). Interestingly, we find that distinct RNAs are enriched in eIF2α-dependent SGs formed after heat shock and in eIF2α-independent SGs induced by hippuristanol treatment. Heat shock enriches for longer RNAs with lower translational efficiency (Fig. 6a,b, Supplementary Fig. 12a-c), similar to the pattern of mRNAs that co-purify with arsenite-induced stress granule cores^55^. In contrast, these factors show little or no correlation with enrichment after hippuristanol treatment (Fig. 6c,d, Supplementary Fig. 12d-f), suggesting that different transcripts may enter granules depending on the nature of the stress.

**Figure 6.**
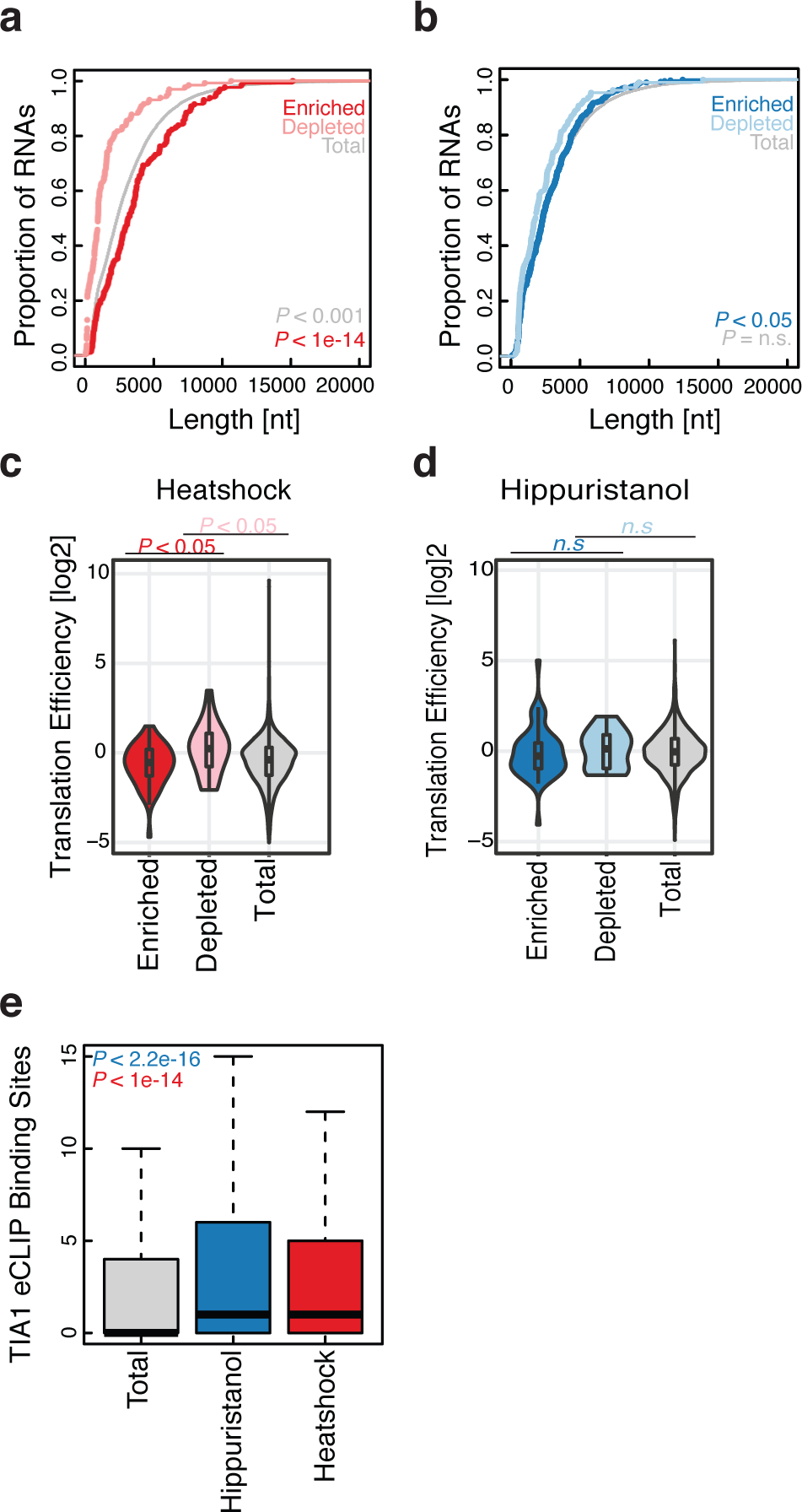
RNA composition of stress granules differs according to the stressor. **a**, Cumulative distribution plot for enriched and depleted RNA length for heat shock stress granule (T_20_ vs T_0_). **b**, As in (**a**) for hippuristanol stress granules. **c**, Translation efficiency distributions for enriched and depleted RNAs during heat shock granule formation. **d**, As in (**c**), for hippuristanol stress granule formation. **e**, TIA1 eCLIP binding site counts for either total, or enriched RNAs during hippuristanol or heat shock stress granule formation. P-values calculated using the Mann-Whitney test. Enrichment and depletion is calculated as statistically significant (FDR <0.05) and either top or bottom 2.5% of the fold change.

Interestingly, while RNAs with longer ORFs were enriched in heat-induced stress granules, they took longer to enter these granules after heat shock. The length bias we saw 20 minutes after heat shock (T_20_) was absent 4 minutes (T_4_) post heat shock (Supplementary Fig. 12g). This change was driven by the accumulation of long transcripts in stress granules at later timepoints: ~45% of the RNAs enriched at T_4_ were also enriched at T_20_; in contrast, only ~12% of T_20_-enriched transcripts were also enriched at T_4_. The slower entry of long RNAs into stress granules may reflect the fact that elongating ribosomes take longer to finish translation on longer CDSes. Stress granule formation requires elongating ribosome to disengage from mRNAs^56^, and run-off elongation from longer ORFs should take more time than run-off from shorter ORFs after stress-induced initiation shutoff.

Key stress granule markers such as TIA1 and G3BP1 are RNA-binding proteins, and indeed, we saw strong enrichment for RBPs in our proteomics data (Supplementary Fig. 8e,10). We thus took advantage of available enhanced RNA crosslinking and immunoprecipitation (eCLIP) data to search for correlations between SG enrichment and protein interactions across the transcriptome. In agreement with previous work, we find that RNAs enriched in either the heat shock or the hippuristanol SG transcriptome contain more TIA1 and HNRNPA1 binding sites than expected by chance^55^ (Fig. 6e, Supplementary Fig. 11c). In contrast, other RBPs, including G3BP1, showed no enrichment for heat shock granules (Supplementary Fig. 12h).

## Discussion

We have developed an approach to study sub-cellular RNA localization by proximity labeling and deep sequencing (Fig. 1). The APEX-Seq approach captures patterns of localization both across (Fig. 2) and within different compartments (Fig. 3). Notably, the 10 - 100 nm range of APEX2 proximity labeling and the very general reactivity of radical-based APEX labeling promise powerful new capabilities for characterizing protein-RNA and protein-DNA structures in cells. It will complement chromatin immunoprecipitation (ChIP) and RNA crosslinking and immunoprecipitation (CLIP) by detecting weak or long-range interactions that are not well captured by crosslinking and circumventing the need for protein immunoprecipitation. Furthermore, the same APEX fusion protein can be used to characterize protein and RNA localization, promising new insights into dynamic and heterogeneous protein-RNA complexes within cells.

We have taken advantage of APEX-Seq in conjunction with proteomics to provide new insights into the organization of translation initiation complexes on active mRNAs and the composition of repressive RNA granules. Our data indicate that eIF4A1 assembles onto the 3’ end of the 43S preinitiation scanning complex, near the mRNA entry site (Fig. 4). To date, the identification of eIF4F with respect to the preinitiation scanning complex remains unknown^44,57^. We also find that eIF4A1 and eIF4E1, another component of eIF4F, show similar patterns of enrichment across the transcriptome despite their involvement in nearly all cap-dependent translation. Our results agree broadly with the transcript-specific effects of modulating eIF4F activity.

We also report the RNA as well as the protein composition of stress granules *in vivo*. During heat shock, we find dynamic changes in the organization of the proteome that reflect the assembly of eIF4A1, along with many other proteins, into stress granules. Previous studies have argued that stable stress granule cores^24^ form first, and a more dynamic outer shell then assembles onto them by liquid-liquid phase separation (LLPS)^45^. In agreement with previous work, we find increased labeling of many stress granule core proteins early during granule formation^45^. Interestingly, however, certain core granule interactions appear to occur independent of stress (Fig. 5), suggesting a model in which stress granule cores reflect at least in part the stabilization or enhancement of pre-existing interactions. We also find that the RNA content of stress granules can vary depending on the nature of the stress, with potential impacts on the transcriptome as well as on protein synthesis (Fig. 6).

Matching both the spatial transcriptome using APEX-Seq, and the spatial proteome using APEX-MS is a particularly powerful approach to better understand the organization of the cell. Critically, with the use of ratiometric comparisons, its use extends to non-membrane bound organelles that remain challenging to work with through more classic techniques, such as RNA granules. This approach addresses a need that cannot easily be met using existing techniques. Protein-RNA crosslinking has revolutionized our understanding of RNA-binding proteins and their targets, but it typically requires direct contact on the ~0.3 nm length scale of a covalent bond. Single-molecule fluorescence *in situ* hybridization is capable of measuring sub-cellular RNA localization, but it is limited by the ~100 nm resolution of optical microscopy and can target only a few, pre-chosen transcripts. The intermediate length scale of APEX-Seq is well suited to address important questions about the organization of transcripts during synthesis, processing, transport, translation, and decay.

## Methods

### General

HEK 293 Flp-In T-Rex cells (Invitrogen) were cultured in DMEM + GlutaMAX (ThermoFisher Scientific, 10566-016) with 10% FBS. APEX fusion plasmids were transfected along with pOG44 by X-tremeGENE 9 (Roche) and selected using 150 µg ml^−1^ of Hygromycin B and 15 µg ml^−1^ of Blasticidin to obtain stable integrants. TMT mass spec reagents (Thermo Fisher Scientific) were used for every quantitative mass spec experiment. Biotin-tyramide was purchased from Iris Biotech (catalog #: LS-3500.1000).

### *in vivo* APEX labeling

Cells were plated in 15 cm dish and cultured for 3 days with 1 µg ml^−1^ of tetracycline. Cells were treated like in^2^. Briefly, Biotin-phenol containing (500 µM final) prewarmed DMEM media was added to cells for 30 minutes prior to the start of the experiment. 1 mM final H_2_O_2_ was added to each dish for a total labeling time of 1 minute (unless stated otherwise). Cells were gently agitated for 1 minute, and quenched with 2X quenching buffer (10 mM Trolox and 20 mM sodium ascorbate in DPBS).

### Protein Purification

BL21 Star (DE3) Escherichia coli cells (Invitrogen) transformed with eIF4A1 (WT) or APEX2-eIF4A1 in a 1.5 L culture were cultivated at 37 °C with 50 µg ml^−1^ kanamycin and then grown at 16 °C overnight with 1 mM IPTG. The cell pellets were resuspended in His buffer (20 mM HEPES-NaOH, pH 7.5, 500 mM NaCl, 10 mM imidazole, 10 mM β-mercaptoethanol) with 0.5% NP-40, sonicated, and centrifuged at 35,000g for 20 min. The supernatant was incubated with 1.5 ml bed volume of Ni-NTA Superflow (Qiagen) for 1 h. The beads were loaded on a gravity column and washed with His buffer containing 1 M NaCl. The proteins were eluted with 50 mM Na-phosphate buffer, pH 7.5, 500 mM NaCl, 100 mM Na_2_SO_4_, 250 mM imidazole. Samples were run through an FPLC HiTrap Heparin HP affinity column (GE Healthcare) with no reducing agent for further purification. Fractions 8-11 were collected. Samples were mixed with 0.25 volumes of 80% glycerol, flash-frozen in liquid nitrogen, and stored at −80°C. All purification steps were performed at 4°C.

### *in vitro* APEX labeling

Recombinant APEX2-eIF4A1 (1.5 µM) was pre-incubated with hemin (4.5 µM) at room temperature for 1 hour. Excess hemin was removed by several rounds of gel filtration through a MicroSpin G-25 column (GE Healthcare). This solution was then combined with biotin-phenol, and a five molar excess of a NanoLuc reporter RNA, in addition to 1 mM ADP and 1 mM MgCl_2_. The samples were then combined with APEX labeling buffer (10 mM Tris, pH 7.5, 150 mM NaCl, and 10% glycerol). The reaction started when peroxide was added to the mix and proceeded for a total of 1 minute *in vitro* labeling. The reactions were then stopped using either TRIzol or oligo binding buffer (Zymo). RNA was extracted using manufacturer’s protocol, and loaded onto a dot-blot apparatus.

### Heat shock and Hippuristanol treatment

Cells were plated onto 10 or 15 cm dishes, and expression was induced for 3 days with 1 µg ml^−1^ of tetracycline. Media was then replaced with fresh media containing biotin-tyramide (500 µM) for 30 minutes. Cells were then placed in a 44°C water bath for better heat transfer during heat shock experiments. APEX labeling was performed immediately after each treatment. For the hippuristanol treatment, cells were treated with biotin-phenol containing media and 1 µM of hippuristanol for 30 min prior to the start of the APEX reaction. Labeling occurred for 30 seconds. Labeling reactions were quenched with 2X quenching buffer (see above) and cells were immediately lysed in TRIzol.

### Streptavidin purification of biotinylated protein

After the APEX labeling reaction, cells were quenched with 2X quenching solution (see above) once, followed by a 1X quenching solution wash. One 1X quenching solution wash was used to resuspend cells, which were then gently pelleted and lysed in 800 µl of lysis buffer containing 1X quenching reagents (1% Triton, 0.1% SDS, 20 mM Tris-HCl pH 7.4, 150 mM NaCl, 5 mM MgCl2, 5 mM trolox, 10 mM sodium ascorbate, and one tablet (per 10 ml) of cOmplete Mini Protease Inhibitor Cocktail). Lysates were clarified by centrifugation for 10 min at 20,000 x g, 4°C. Streptavidin beads (Pierce) were equilibrated with lysis buffer for a total of two washes. Lysate was mixed with streptavidin beads at a volumetric ratio of 8 volumes beads per 5 volumes lysate and incubated at RT for 1 hr. Beads were washed twice with lysis buffer, once with 1M KCl solution, once with 2M urea, pH 8, and twice with lysis buffer (w/o detergent or quenching reagents). Biotinylated proteins were eluted using by boiling samples in 8M urea, pH 8 for 3 minutes at 98°C.

### Streptavidin purification of biotinylated RNA

After an *in vivo* labeling reaction, cells were lysed with TRIzol and RNA was purified by by precipitating RNA from the aqueous phase. Samples were treated with DNase I (NEB) and ~100 µg of RNA was then gently fragmented using 10^-5^ units of RNase A/T1 (Thermo Fisher Scientific) by placing 1 µl of the RNase cocktail in the lid of each tube and initiating the reaction by simultaneously centrifuging the samples. These were then incubated for 10 minutes at 37°C in a final RNA concentration of 1µg µl^-1^. The reaction was immediately quenched using 400 µl of TRIzol. RNA was extracted by either adding 500 µl of TRIzol, followed by 200 µl of Chloroform, and by precipitating RNA from the aqueous phase, or by Direct-Zol (Zymo) purifications following the manufacturer’s instructions. C1 Streptavidin beads (10 µl per sample; Thermo Fisher Scientific) were washed three times with Buffer 1 (1 mM MgCl_2_, 0.5% sodium deoxycholate in PBS), washed once and blocked for 30 minutes with Blocking Buffer (5X Denhardt’s reagent and 150 µg/ml Poly IC (InvivoGen) in Buffer 1). Blocking Buffer was removed and replaced with extracted RNA samples in fresh Blocking Buffer. Samples were incubated at room temperature for 1 hour and then washed twice with Buffer 2 (6M Urea, pH 8, 0.1% SDS in PBS), once with Buffer 3 (2% SDS in PBS), once with Buffer 4 (750 mM NaCl, 0.5% sodium deoxycholate, 0.1% SDS), and once with Buffer 5 (150 mM NaCl, 0.5% sodium deoxycholate, 0.1% SDS). RNAs were eluted from Streptavidin beads by denaturation in 300 µl TRIzol (Qiagen) and extracted using either the above mentioned TRIzol/Chloroform procedure or Direct-zol kit, following the manufacturer’s instructions. RNAs were eluted in 6 µl H_2_O. RNAs were then fragmented for 7 minutes (as recommended for RIN values > 2 and < 7) at 94°C, following the NEBNext Ultra II Directional RNA Library Prep Kit (NEB, catalog # E7760S) instructions.

### TMT mass spec

100 µg of eluted proteins were brought up to 100 µl with 100 mM TEAB. 5 µl of 200 mM TCEP were added and samples were incubated at 55°C for 1 hour. Fresh iodoacetamide was made up in 100 mM TEAB (final concentration 375 mM). 5 µl of the 375 mM iodoacetamide were added to the sample and incubated for 30 minutes protected from light at room temperature. Six volumes (or more) (~600µL) of pre-chilled (−20°C) acetone were added. Samples were allowed to precipitate overnight at −20°C. Samples were precipitated at 8,000 x g for 10 min at 4 °C and resuspended with 100 µl of 50 mM TEAB. For the tryptic digest, 2.5 µl of trypsin (1 µg/µl) were added to 100 µg of protein and incubated overnight at 37 °C. Peptides were quantified and normalized using a quantitative colorimetric peptide assay (Thermo Fisher Scientific). 41 µl of anhydrous acetonitrile were added to each TMT labeling reagent vial and the reagent was allowed to resuspend at RT for 5 min with occasional vortexing. The reduced and alkylated protein digest were transferred to each TMT Reagent vial. Reactions were incubated for 1 hr at RT. 8 µl of 5% hydroxylamine was added to each sample and incubated for 15 minutes to quench the reaction. Samples were dried in a Speed-Vac or frozen in −80°C for storage and transportation. Samples were run on an OrbiTrap mass spectrometer (ThermoFisher Scientific) using high pH fractionation, and analyzed using Proteome Discoverer. DESeq2^58^ was used to perform quantitative ratiometric comparisons between each APEX fusion and APEX2-GFP.

### Western Blots

Samples were blocked with 0.6% milk in TBST (0.05% tween-20) for 1 hour at RT. Membranes were incubated with the corresponding primary antibodies in signal enhancer solution 1 (Hikari NU00101) for 1 hour at RT, and secondary antibodies in signal enhancer solution 2 (Hikari NU00102) for 1 hour at RT. Three 5 min washes were performed between antibody incubations. STREP-HRP (1:1,000, CST 3999) was used to blot for biotinylated protein. Anti-FLAG (1:1,000, CST 2368), and anti-eIF4A (1:1,000, CST 2490) antibodies were used as primary antibodies and HRP-conjugated anti-rabbit IgG (CST 7074) was used as a secondary antibody. Chemiluminescence was induced by SuperSignal West Dura Extended Duration Substrate (Thermo Scientific) and images were acquired by a FluorChem R imaging system (ProteinSimple).

### Immunofluorescence

Cells were plated onto poly-lysine coated coverslips in 12 well plates and APEX fusion expression was induced by adding 1 µg ml^-1^ of tetracycline. Two days later, cells were fixed in 3.7% paraformaldehyde (PFA) in PBS for 10 minutes at room temperature, and permeabilized with 1% Triton X-100 in PBS for 10 min at room temperature. Cells were then blocked with 3% BSA in PBS for 30 min at room temperature. All primary antibodies were incubated overnight at 4 °C and secondary antibodies at 37 °C for 1 hour. Several PBS washes were performed in-between antibody staining. Coverslips were mounted onto slides with ProLong Gold mountant (Thermo Fisher Scientific).

### Data and code availability

Raw sequencing data are available for download GEO: GSE121575. Scripts to run the analyses mentioned above are available upon request.

### Dot Blot Assay

All dot blot assays were performed using the Dot-Blot Microfiltration Apparatus (Bio-Rad). Zeta-Probe membranes (Bio-Rad) were submerged in H_2_O for 5 min. RNA samples were loaded onto the apparatus and the solution was gently pulled through the membrane by vacuum suction (setting 3). Zeta-probe membranes were then cross-linked two times at 200 µJ/cm^2^ (CL-1000 Ultraviolet Crosslinker) and blocked for 2 hr using Odyssey blocking buffer + 1% SDS. 800CW Streptavidin (1:10,000 LI-COR Biosciences) was then added for 30 minutes in fresh blocking buffer with 1% SDS. Membranes were visualized in the LI-COR (Odyssey CLx). A ssDNA oligo with a 3’-biotin modification was used as a positive control (IDT).

### Classifier predicting RNA granule residency

A regularized logistic regression classifier was built using biophysical features from each protein and known protein domains from PFAM. Continuous features were scaled and centered. 70% of the data was partitioned into a training set. Repeated k-folds cross validation (k = 10) was performed for alpha and lambda parameters that minimized mean squared error.

## ACKNOWLEDGEMENTS

We thank members of the Ingolia and Lareau labs as well as A. Dernberg, and L.J. Kitch for discussion. This work was supported by an NIH New Innovator’s Award DP2 CA195768-01 (N.T.I.), an NSF Graduate Research Fellowship (A.P.), and a Human Frontiers in Science Program long-term fellowship (S.I.). This work used the Vincent J. Coates Genomics Sequencing Laboratory at UC Berkeley, supported by NIH Instrumentation Grants S10 RR029668, S10 RR027303, and S10 OD018174. Proteomics data was acquired at the UC Davis Genome Center Proteomics Core by M. Salemi and B. Phinney.

## AUTHOR CONTRIBUTIONS

A.P. and N.T.I. conceived and designed the study; A.P. and S.I. performed the experiments; A.P. and N.T.I. wrote the manuscript with contributions from all authors.

## COMPETING INTERESTS

The authors declare no competing financial interests.

